# Brain-body mitochondrial distribution patterns lack coherence and point to tissue-specific and individualized regulatory mechanisms

**DOI:** 10.1101/2024.09.20.614152

**Authors:** Jack Devine, Anna S Monzel, David Shire, Ayelet M Rosenberg, Alex Junker, Alan A Cohen, Martin Picard

## Abstract

Energy transformation capacity is generally assumed to be a coherent individual trait driven by genetic and environmental factors. This predicts that some individuals should have high and others low mitochondrial oxidative phosphorylation (OxPhos) capacity across organ systems. Here, we test this assumption using multi-tissue molecular and enzymatic activities in mice and humans. Across up to 22 mouse tissues, neither mitochondrial OxPhos capacity nor mtDNA density were correlated between tissues (median *r* = -0.01–0.16), indicating that animals with high mitochondrial capacity in one tissue can have low capacity in other tissues. Similarly, the multi-tissue correlation structure of RNAseq-based indices of mitochondrial gene expression across 45 tissues from 948 women and men (GTEx) showed small to moderate coherence between only some tissues (regions of the same brain), but not between brain-body tissue pairs in the same person (median *r* = 0.01). Mitochondrial DNA copy number (mtDNAcn) also lacked coherence across organs and tissues. Mechanistically, tissue-specific differences in mitochondrial gene expression were attributable in part to i) tissue-specific activation of canonical energy sensing pathways including the transcriptional coactivator PGC-111 and the integrated stress response (ISR), and ii) proliferative activity across tissues. Finally, we identify subgroups of individuals with high mitochondrial gene expression in some tissues (e.g., heart) but low expression in others (e.g., skeletal muscle) who display different clinical phenotypic patterns. Taken together, these data raise the possibility that tissue-specific energy sensing pathways may contribute to the idiosyncratic mitochondrial distribution patterns associated with the inter-organ heterogeneity and phenotypic diversity among individuals.

## Introduction

A major driver of organ-specific function and dysfunction is the capacity for energy transformation. Energy enables organ-specific function and inter-organ communication ^1,2^. In breathing animals, energy is transformed within mitochondria, where the oxidative phosphorylation (OxPhos) system converts oxygen and food substrates into usable cellular energy. Because we develop from a single fertilized mitochondria-filled oocyte into a mature adult composed of genetically identical cells and mitochondria, it is generally assumed that inherited (epi)genetic factors define mitochondrial OxPhos capacity and biology homogenously across the whole body. In single-tissue studies such as blood immune cells, genetic variants explain a portion of the variance in mitochondrial DNA copy number (mtDNAcn) ^3^. Similarly, exercise can induce biogenesis not only in the working muscles but also in the brain and other tissues ^4,5^, suggesting that behaviorally-driven mitochondrial adaptations occur systemically across the whole body, via the action of only partially resolved factors ^6^. Therefore, based on these premises, there is an expectation that organisms should display mitochondrial inter-tissue “coherence”. This means that relative to other individuals in the population, an individual who has high mitochondrial content in one tissue (e.g., brain) would also have high mitochondrial content in other tissues (e.g. heart, skeletal muscle, skin, etc.). Similarly, we would expect some individuals to have low mitochondrial content across all tissues (**Figure 1A**).

**Figure 1.**
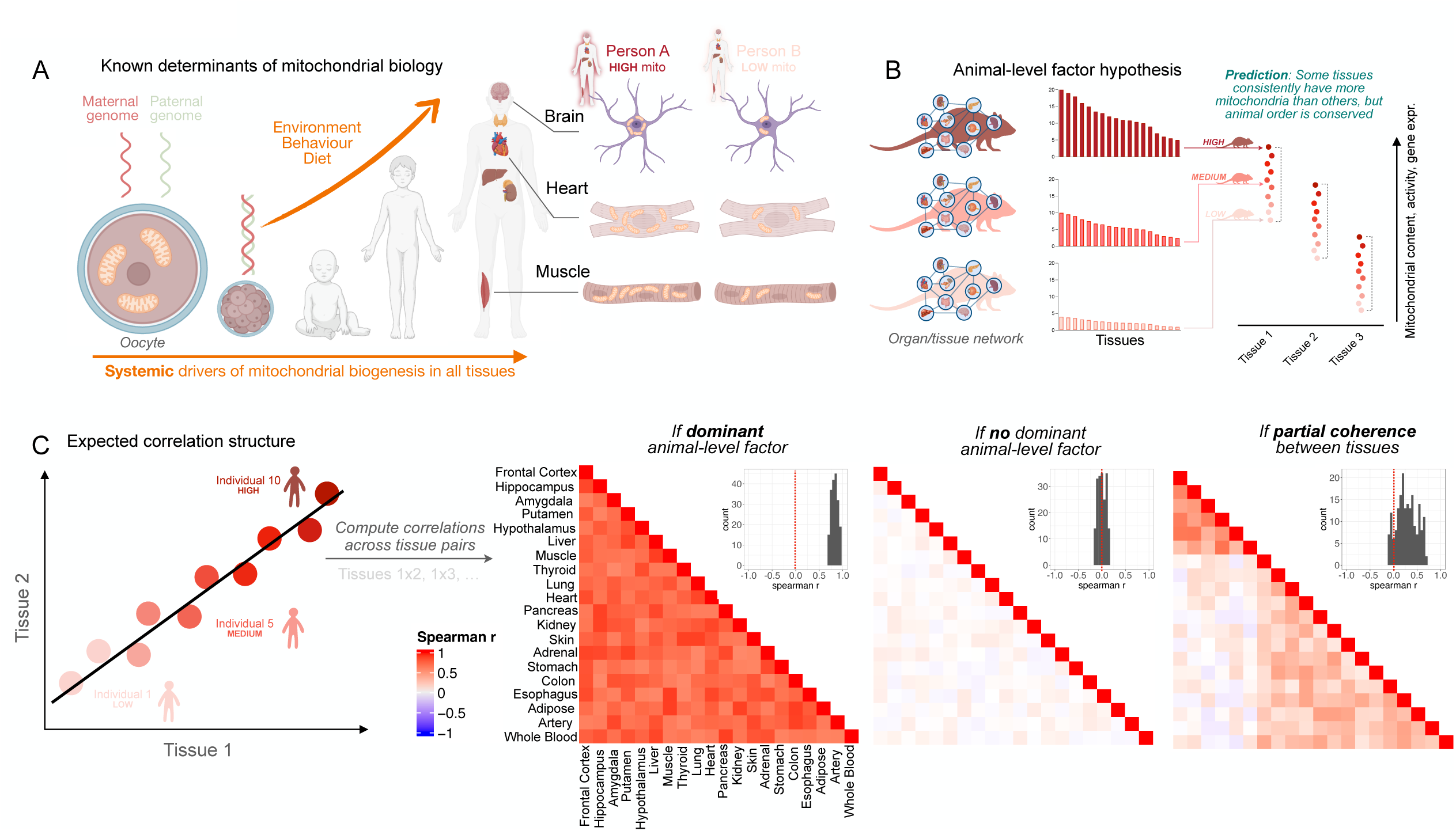
Expectations and hypotheses on individual phenotypes of multi-tissue mitochondrial biology. Figure illustrating the multi-tissue coherence hypothesis. The expectation is that some individuals will have high mitochondrial content in all tissues and some will have low mitochondrial content in all tissues. This would result in strong correlations displayed between tissues in measures of mitochondrial biology.

If the inter-tissue coherence hypothesis is true, measurements of mitochondrial OxPhos capacity and gene expression across multiple tissues would be expected to resemble **Figure 1B**. Tissues with high baseline energetic demand (e.g. heart) consistently have more mitochondria than others with lower demand. But most importantly, the animal order should be conserved across tissues: some animals should exhibit high values in all tissues, and some animals should be low in all tissues. The inter-tissue correlation structure observed from this result would be strongly positive, displaying coherence between tissues (**Figure 1C**). However, if a different correlation structure is observed between tissues (weak, absent, or negative correlations) this would suggest that the individual-level, inter-tissue coherence hypothesis is incorrect. The assumption of inter-tissue coherence underlies much biological research, although rarely made explicit and seldom tested, with a few exceptions. Notably, the lack of coherence in mtDNAcn and mitochondrial respiration across human tissues ^7,8^ together with recent evidence that different human organs age at different rates relative to one another even in a given body ^9–11^ brings the inter-tissue mitochondrial coherence hypothesis into question.

Here we systematically test this hypothesis at scale using multi-tissue OxPhos enzymatic activity measurements in two independent mouse cohorts, plus in whole transcriptomes from 45 tissues from 948 women and men (n=16,205 samples, n=983 total tissue-tissue pairs). Unlike the default hypothesis that mitochondrial biology is predominantly an organism-level attribute of each individual, our results highlight the general lack of coherence across brain and non-brain organ systems. These results point towards the emergence of idiosyncratic mitochondrial distribution patterns across individuals, potentially driven by genetic regulatory pathways associated with organ-specific mitochondrial gene expression patterns.

## Results

### Multi-tissue mitochondrial enzymatic activities and mtDNA density display low between-tissue correlations in two independent mouse cohorts

We first tested the trait-level inter-tissue coherence hypothesis using direct biochemical enzymatic activities for OxPhos complexes I, II and IV, and the Krebs cycle enzyme citrate synthase (CS) in 5 tissues: Brain – hippocampus; non-brain – liver, brown fat, muscle and bone from 16 male mice. mtDNA density was also quantified in each tissue by qPCR (n=10 tissue pairs x 5 measures per tissue giving a total of n=50 pairwise comparisons for specific mitochondrial features compared between tissues). Contrary to the hypothesis, the resulting correlation matrix of the 5 mitochondrial measures across the five tissues showed minimal or even negative correlation between tissues (**Supplemental Figure 1**). The frequency distribution of the correlation coefficients (Spearman r) across the 50 tissue pairs is shown in **Supplemental Figure 1B**. Individual biplots in which each data point represents a different animal show the small-to-null correlations across organ systems.

Intrigued by the consistency of these findings across multiple mitochondrial enzymes from several tissues, we replicated this analysis in a second cohort of 27 male mice in which we quantified OxPhos enzyme activities and mtDNA density in 17 brain regions and 5 non-brain tissues (n=1,155 tissue-tissue pairs), the largest multi-tissue mitochondrial biochemistry dataset available to our knowledge. The resulting correlation matrix containing all measures across all tissues shown in **Figure 2A** is divided into brain-brain, brain-body, and body-body correlations (see inset **Figure 2A**). Small (r=0.1-0.3), moderate (r=0.3-0.5) and some large (r>0.5) correlations were observed between brain regions, reflecting a certain degree of coherence between different parts of the same organ (median r=0.25). Of the brain-brain correlations, 86% were positive (19.7% significant, uncorrected p<0.05) and 14% were negative (0.15% significant, uncorrected p<0.05). In contrast, there was no coherence between brain-body tissues (median r=0.03) and between various non-brain tissues (r=-0.03) (**Figure 2B**). In fact, some correlations were even negative: animals with high amygdala complex IV activity tended to have lower adrenal gland complex IV activity (r=-0.54, uncorrected p<0.01, **Figure 2F**). This could indicate a potential functional tradeoff between some tissues, where organisms who have high mitochondrial expression in one tissue have low expression in another (discussed further below).

**Figure 2.**
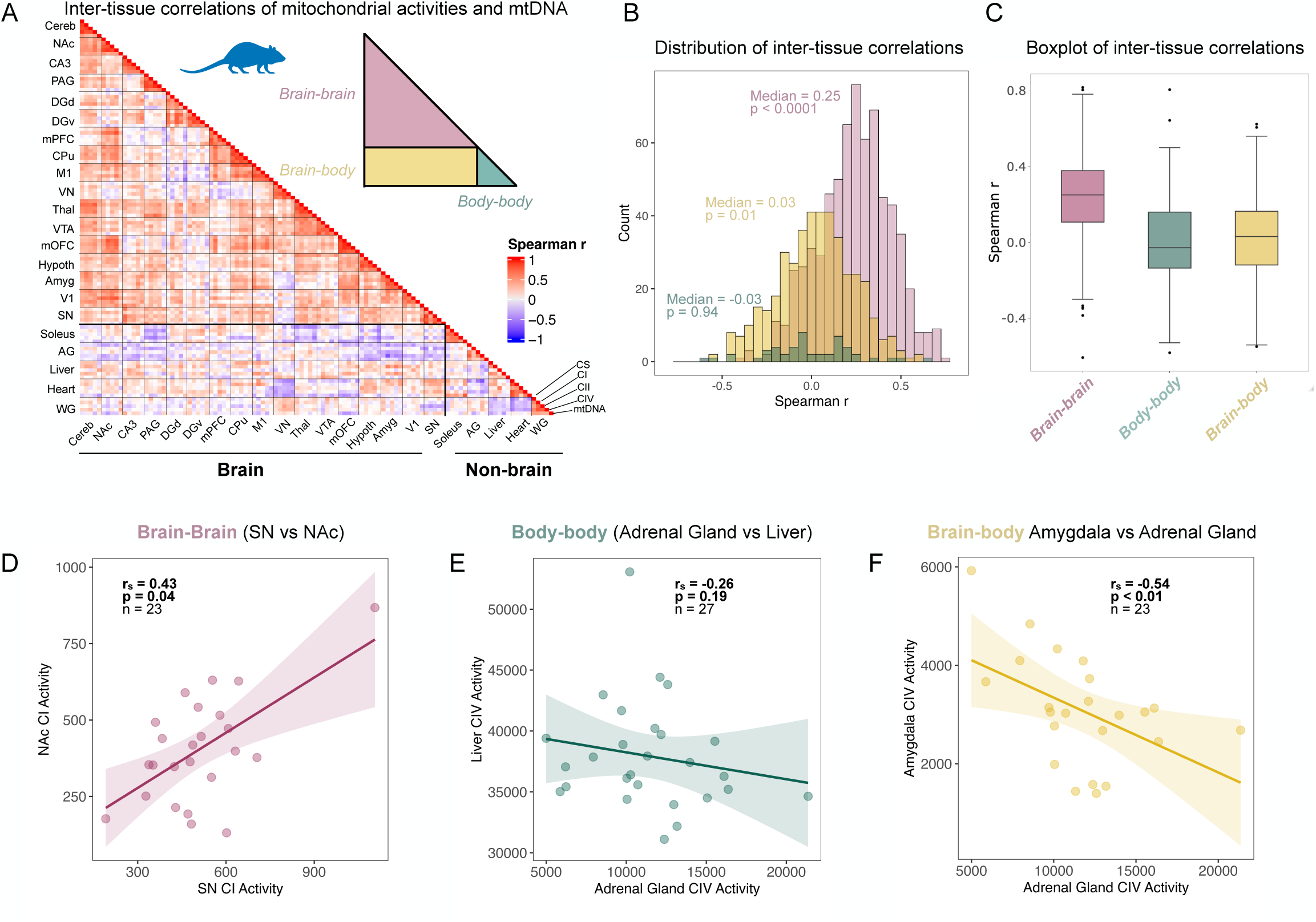
Mitochondrial enzymatic activity and mtDNA density measures display low between tissue correlations across 22 tissues from 27 mice. **(A)** Enzymatic activity and mtDNA density-based multi-tissue mitochondrial distribution patterns displayed as correlation matrix of enzymatic activity measures (CI, CII, CIV and CS) and mtDNA density across 17 brain tissues and 5 peripheral tissues from 27 male mice in Cohort 2 (n=1,155 tissue pairs). (B) Frequency distribution of spearman r correlation coefficients of pairwise tissue comparisons showing brain-brain (red), body-body (green), and brain-body (yellow) tissue comparisons. (C) Boxplot of brain-brain, body-body and brain-body inter-tissue correlations. (D-F) Bivariate plots of mitochondrial enzyme activity measures between tissues**. Abbreviations:** Cereb = Cerebellum, NAc = Nucleus Accumbens, CA3 = CA3 region, PAG = Periaqueductal grey, DGd = dorsal dentate gyrus, DGv = ventral dentate gyrus, mPFC = medial prefrontal cortex, CPu = caudoputamen, M1 = primary motor cortex,, VN = vestibular nucleus, Thal = thalamus, VTA = ventral tegmental area, mOFC = medial orbitofrontal cortex, Hypo = hypothalamus, Amyg = amygdala, V1 = primary visual cortex, SN = substantia nigra, Soleus = red oxidative skeletal muscle, AG = adrenal gland, WG = white glycolytic skeletal muscle.

Nevertheless, contrary to the default hypothesis that correlations would be positive and indicate inter-tissue coherence, 93.4% of brain-body correlations were not statistically different from 0 (45% negative, 55% positive; only 3.3% were significantly positive, and 3.3% were significantly negative, uncorrected p<0.05). The same was observed between non-brain tissues, where 84% of the tissue-tissue correlations were not statistically different from 0 (54% negative, 46% positive; 8% significantly negative, and 8% significantly positive), roughly indicating chance-level results.

Thus, animals with high OxPhos capacity and mtDNA content in one tissue do *not* tend to also have high OxPhos capacity in other tissues, demonstrating an overall lack of coherence in mitochondrial biology between tissues of the same organism. Both mouse datasets are available in **Supplemental File 1**.

### Transcriptome-based mitochondrial profiling

Next, we examined the trait-level inter-tissue coherence hypothesis in humans. We leveraged the Genotype Tissue Expression (GTEx) ^12^ RNAseq data from 948 women and men across 45 tissues (16,205 samples), which similarly allows to perform multiple brain-brain, brain-body, and body-body tissue comparisons. Using gene expression data for each tissue we quantified the percentage of all mRNA transcripts that are derived from the mitochondrial genome (mtDNA%), reflecting most directly, albeit imperfectly, the mass or content of mitochondria within each person/tissue. Separately, based on the inventory of mitochondrial genes Mitocarta 3.0 ^13^, we quantified for each GTEx participant and tissue the proportion of the nuclear transcriptome (all nuclear transcripts arising from ∼20,000 genes) that is “invested” to produce the ∼1,100 mitochondrial proteins (mito-nDNA%). This yielded novel quantitative indices of mitochondrial regulation for each participant-tissue combination (**Figure 3**), available in **Supplemental File 2**.

**Figure 3.**
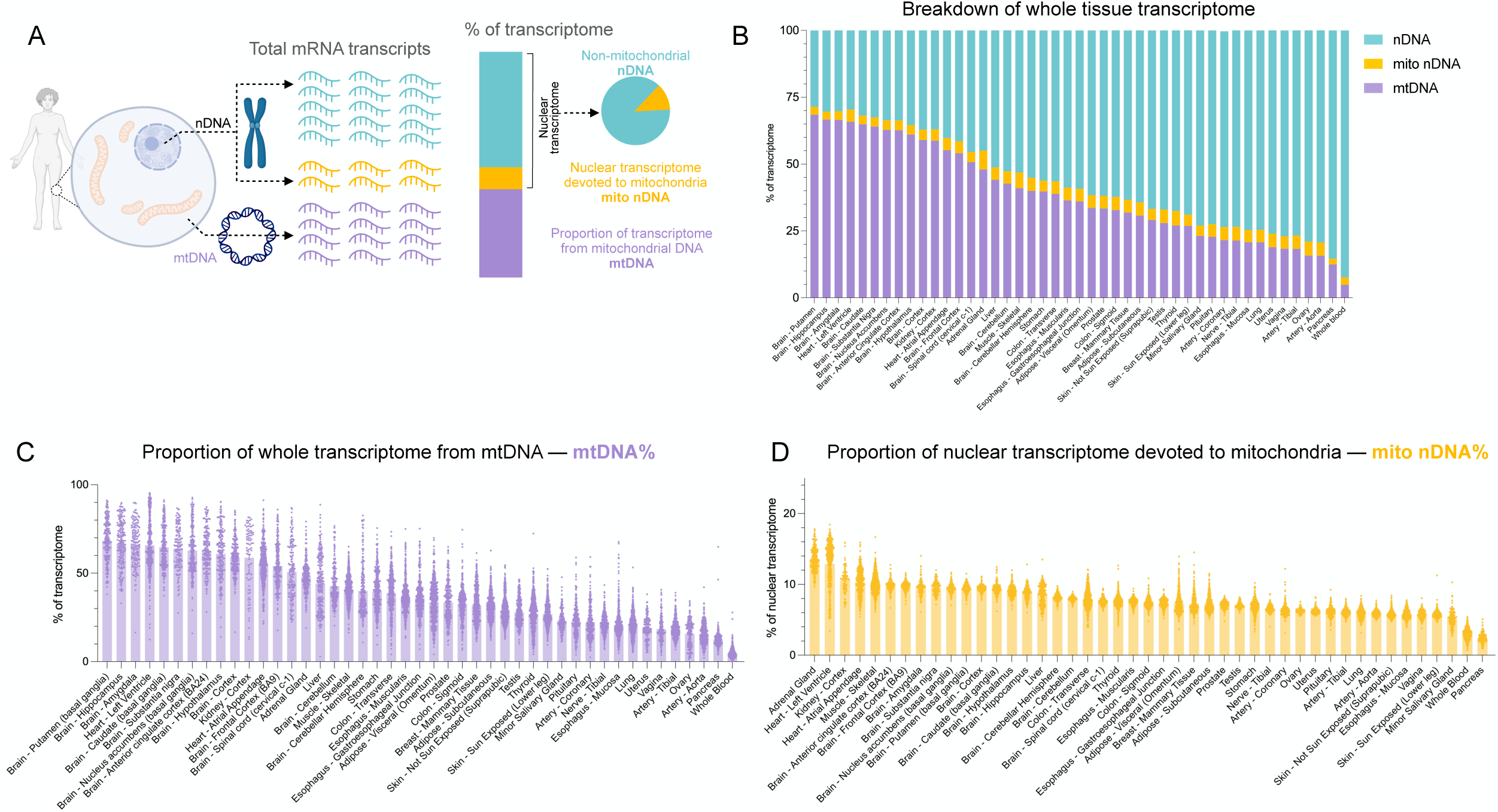
Transcriptome-based mitochondrial profiling across human tissues. (A) The percentage of total transcripts from the mtDNA, nuclear mitochondrial genes and non-mitochondrial nuclear genes were quantified in each sample (n=16,205). (B) The average percentage of transcripts that are from mtDNA and nuclear mitochondrial genes (1120 genes Mitocarta 3.0) in each tissue. (C) Ranked mean percentage of total transcripts that are mtDNA transcripts in each tissue (each datapoint represents a person). (D) Ranked mean percentage of nuclear transcripts that are transcripts of mitochondrial nuclear genes (each datapoint represents a person).

As expected, the tissues with the highest fraction of mtDNA-derived transcripts (mtDNA%) were brain regions (putamen, hippocampus, amygdala), followed by the heart (left ventricle), with mtDNA transcripts composing on average ∼60-70% of all mRNA transcripts (**Figure 3B**). In a few individual participant’s brains and hearts, >90% of mRNAs were of mtDNA origin. The lowest ranking tissues in mtDNA% were pancreas and whole blood, where only ∼4.8-12.5% on average of the cellular transcriptome is from the mitochondrial genome. This result was expected as blood leukocytes, particularly the abundant neutrophils, have few mitochondria and few mtDNA copies per cell ^14^, and participants had on average 4.8% mtDNA% (range 0.4%-27.8%). The difference between the average of the highest and lowest ranking tissue was 14.1-fold, reflecting the well-known natural variation in mtDNA copies and mitochondrial abundance between human tissues ^7,15^.

The tissues with the highest proportion of *nuclear* transcripts devoted to mitochondria (mito-nDNA%) were adrenal gland and heart (left ventricle), consistent with their high mitochondrial volume density ^16,17^. As in mtDNA-derived transcripts, the lowest ranking tissues for nuclear transcripts were whole blood and pancreas (**Figure 3D**). As expected from well-established and stable tissue differences in mitochondrial content, tissues with higher mtDNA% also generally express higher mito-nDNA% (r=0.84, p<0.0001, **Supplemental Figure 2**), lending validity to these measures. We analyze the association of these parameters at the individual level within each tissue below.

### Human mitochondrial gene expression patterns display weak brain-body coherence

Having established proximate markers of mitochondrial abundance and expression from RNAseq data, we then tested the inter-tissue coherence hypothesis in humans by quantifying mtDNA% and mito-nDNA% for all available GTEx subjects across 45 organs and tissues. We proceeded to compute the correlations across all tissue pairs (n=983 tissue-tissue pairs) and analyzed the multi-tissue correlation structure (**Figure 4A**). A heatmap representing the multi-tissue correlation structure of mito-nDNA% is shown in **Figure 4B**, divided into brain-brain, body-body, and brain-body comparisons.

**Figure 4.**
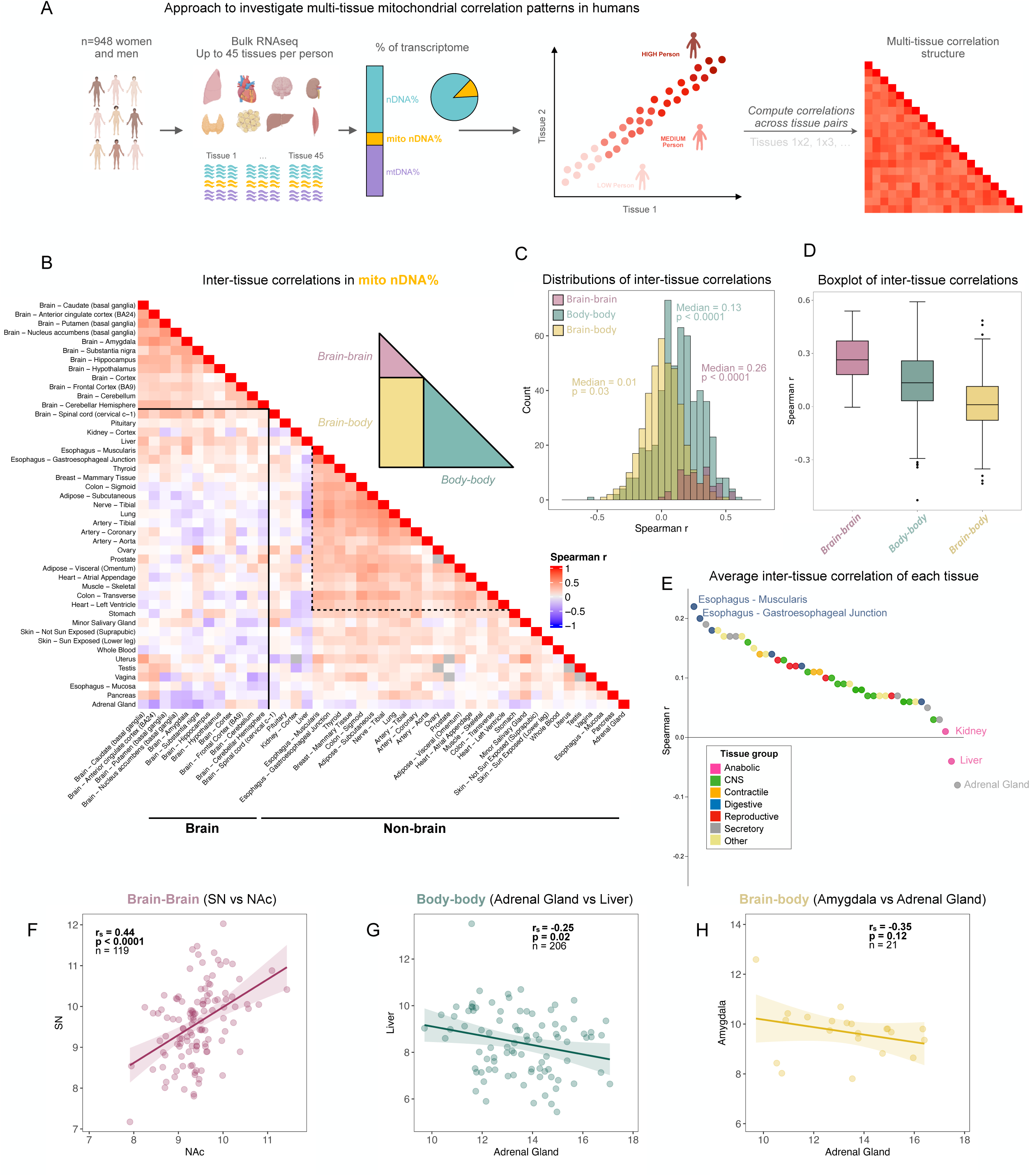
Mitochondrial gene expression patterns in human tissue display low between-tissue correlations. (A) Experimental set-up of multi-tissue correlation analysis of GTEx RNAseq data. (B) Heatmap of correlation matrix showing the pairwise spearman r correlations of mito-nDNA% between 45 organs/tissues. The dotted line is highlighting body tissues showing medium positive correlations. (C) Frequency distribution of spearman r correlation coefficients (983 tissue comparisons) between brain-brain, body-body and brain-body tissues. (D) Boxplot displaying median of spearman r correlation coefficients of brain-brain, body-body and brain-body tissue pairs. (E) Average inter-tissue correlation of each tissue. (F-H) Bivariate plots of mito-nDNA% between tissues.

The highest between tissue correlations were observed between the 12 brain areas with a median of r=0.26 (**Figure 4B**). This result is similar to the median correlation (r=0.25) observed between the 17 brain areas in mice (**Figure 2B**). Correlations of body tissues with other body tissues had a median of r=0.13. As in mouse tissues, several human non-brain tissues showed medium positive correlation indicating some coherence between specific tissues (highlighted with dotted line, **Figure 4B**), while others showed weak positive, or even negative correlations. Again, the brain-body tissue pairs displayed minimal correlations (median r=0.01, **Figures 4C-D**). The adrenal gland exhibited the lowest degree of coherence with other tissues (mean r=-0.08), while the esophagus was most the coherent with all other tissues (mean r=0.22) (**Figure 4E**). As in mice, the brain substantia nigra and nucleus accumbens showed medium positive correlation (r=0.44, p<0.0001, **Figure 4F**), consistent with conserved cross-species neuroanatomical and functional connectivity. Also in agreement with our mouse cohort, the amygdala and the liver both showed negative correlations with adrenal gland (**Figure 4G-H**).

Individuals with greater mito-nDNA% in their amygdala and liver tended to have lower adrenal gland mito-nDNA%, indicating minimal co-regulation and potentially the existence of negative tradeoffs between these tissues.

Repeating this analysis with mtDNA% of transcriptome, or with specific functional mitochondrial pathways, yielded similar results (**Supplemental Figure 4**). While different mitochondrial pathways (specialized subsets of genes) exhibited relatively distinct correlation structures, all exhibited medium correlations between brain regions and among some body tissues. However, regardless of the mitochondrial gene subsets selected, there was a uniform lack of coherence among brain-body tissue pairs, confirming the results above for mito-nDNA%. The multi-tissue network architecture of mitochondrial gene expression highlighting tissue pairs with the strongest and weakest coherence are visualized in **Figure 5**. The equivalent network for mtDNA% is presented in **Supplemental Figure 5**. Tissues of the same type (e.g., brain regions, digestive tube segments) exhibit the greatest coherence, as expected, highlighting the tissue specialization of mitochondria ^18^.

**Figure 5.**
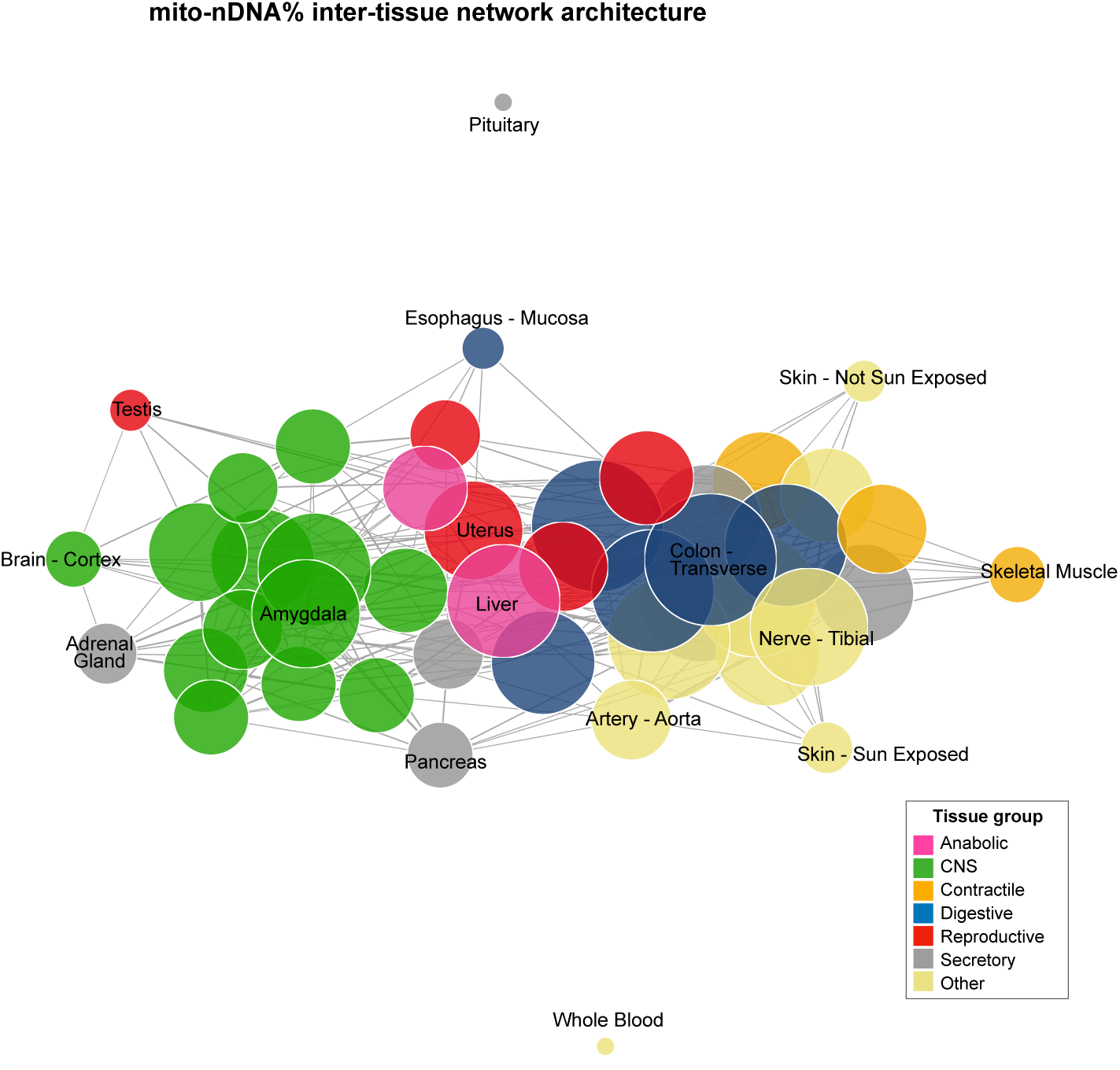
Multi-tissue network architecture of mitochondrial gene expression. Network representation of multi-tissue correlation matrix of mito-nDNA%. Each node represents a tissue, the size of each node is proportional its degree, edge thickness is proportional to the strength of the correlation.

### mtDNAcn-based inter-tissue correlation patterns are consistent with mitochondrial transcript-based patterns in humans

To validate our transcriptome-based findings indicating low coherence between brain and body tissues using a different method, we leveraged the recently reported qPCR-based catalog of mtDNAcn across 52 tissues from 952 GTEx subjects ^15^. We asked if individuals who have higher or lower mtDNAcn relative to other people in one tissue also have higher or lower mtDNAcn in other tissues. The resulting inter-tissue correlation structure of mtDNAcn across the same 45 tissues used in our transcript-based analyses is shown in **Supplemental Figure 6**.

The resulting correlation pattern was highly consistent with that observed using transcript-based indices of mitochondrial expression. Again, brain-body tissue pairs exhibited the lowest coherence (median r=-0.03). Body-body inter-tissue correlations had a median of r=0.10. And again, the highest coherence was observed between brain tissues (median r=0.24), with an effect size highly consistent with both the mouse (median r=0.25) and human transcript-based indices of coherence (median r=0.26) Thus, these results confirm the consistent yet modest coherence between regions of the same organ (brain) and the lack of coherence in mitochondrial biology across brain and body tissues.

### Low inter-tissue coherence at the proteomic level

We also tested the trait-level inter-tissue coherence hypothesis at the level of proteins using the same approach as above but with available proteomics data from a subset of the GTEx cohort: 201 samples from 32 tissues across 14 GTEx subjects (n=19 tissue-tissue pairs after filtering the data) ^19^. There was an insufficient number of brain samples in this dataset to perform brain-body inter-tissue correlation analyses. The resulting multi-tissue correlation structure and frequency distribution of correlation coefficients (**Supplemental Figure 7**) agreed with the transcriptomics data, indicating no strong inter-tissue coherence. Several tissue pairs showed weak, and in some cases negative, correlations in mitochondrial protein abundance, meaning that an individual with high mitochondrial protein abundance in one tissue can have low abundance in others. Although the strength of the correlations are likely inflated due to the low sample size in this analysis, the median correlation coefficient (r=0.26) from the proteomic data in non-brain tissues indicated potential modest inter-tissue coherence.

### Mitochondrial gene expression is driven in part by canonical energy sensing pathways

If mitochondrial content is not driven by a systemic trait-level factor, this raises the question: *what determines the abundance of mitochondria in each tissue of an organism?* We hypothesized that mitochondrial content may be driven by two potential pathways: 1) the transcriptional co-activator and master regulator of mitochondrial biogenesis peroxisome proliferator-activated receptor gamma coactivator 1-alpha (PGC-1α) ^20^, or 2) the integrated stress response (ISR), which is a pathway that regulates energy balance and is induced by reductive stress and mitochondrial OxPhos defects in a tissue-specific manner ^21,22^. If correct, PGC-1α and ISR expression should exhibit positive intra-tissue correlations with mitochondrial gene expression.

Expression of PGC-1α was quantified by calculating the percentage of nuclear transcripts mapping to PGC-1α, and then correlating this to mito-nDNA% (**Figure 6A**) or mtDNA% (**Supplemental Figure 8A**), separately for each of the 45 tissues. As expected in skeletal muscle and heart where PGC-1α was initially discovered to induce mitochondrial biogenesis ^23,24^, individuals with high PGC-1α expression had significantly higher values for mito-nDNA%. This pattern was observed in 24 tissues (ps<0.05). The highest correlations were observed for the transverse colon and left ventricle of the heart. However, 7 tissues exhibited significant negative correlations (ps<0.05), including the adrenal gland, visceral and subcutaneous adipose tissues where greater PGC-1α expression was related to lower mitochondrial gene expression (rs=-0.24 to -0.36, ps<0.05).

**Figure 6.**
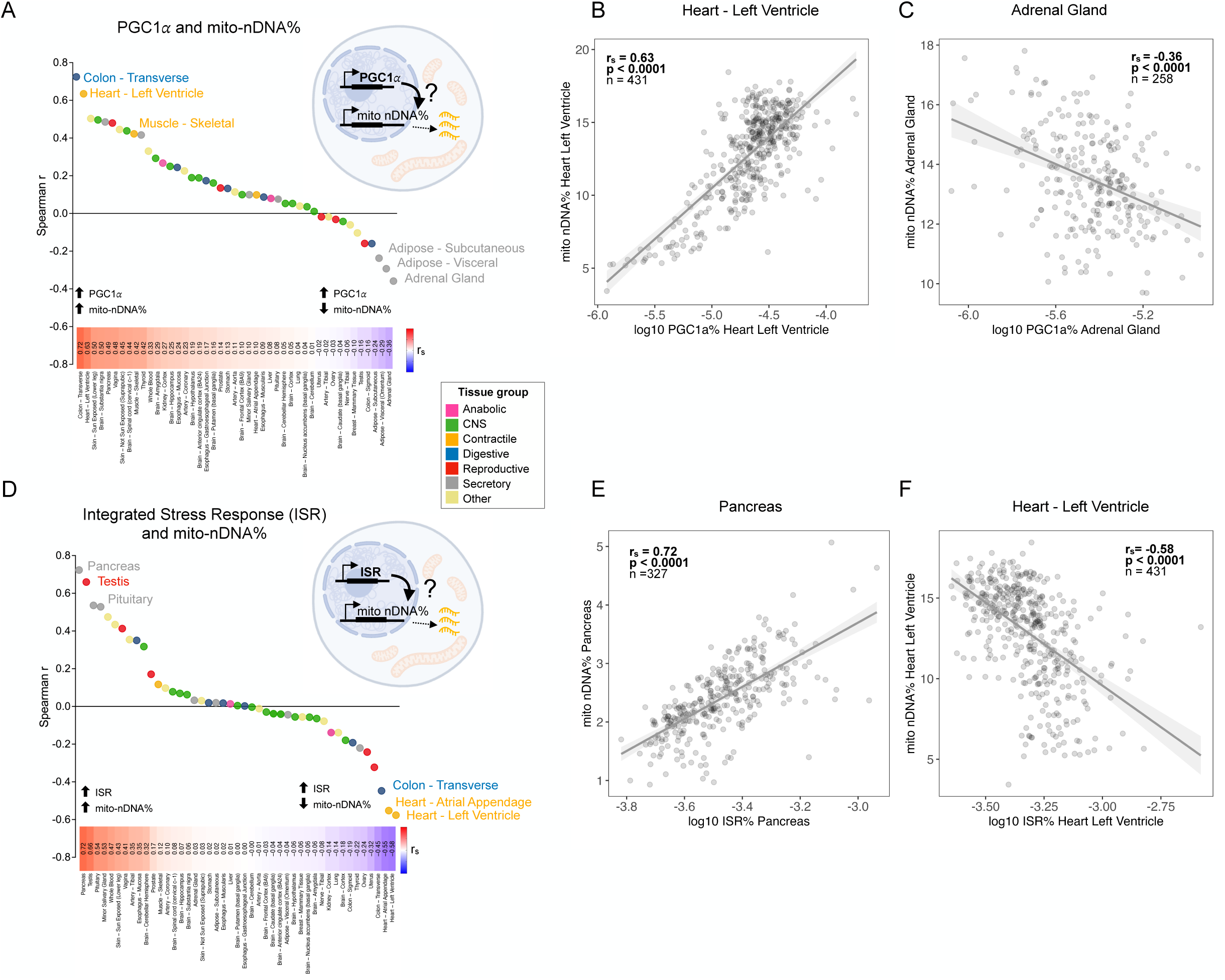
Mitochondrial gene expression is driven in part by canonical energy and stress sensing metabolic pathways. (A) Ranked spearman r correlation coefficients of PGC-1α% vs mito-nDNA% transcripts in each tissue. (B) Biplot showing the correlation of mito-nDNA% with PGC-1α expression in the heart (left ventricle). (C) Biplot showing the correlation of mito-nDNA% with PGC-1α expression in the adrenal gland. (D) Ranked spearman r correlation coefficents of ISR% vs mito-nDNA% transcripts in each tissue. (E) Biplot showing the correlation of mito-nDNA% with ISR expression in the pancreas. (F) Biplot showing the correlation of mito-nDNA% with ISR expression in the heart (left ventricle).

The ISR was quantified by calculating the percentage of nuclear transcripts mapping to four key mammalian ISR genes: ATF4, ATF5, DDIT3 and GDF15 ^25–28^. Like PGC-1α, ISR expression was significantly associated with higher mito-nDNA%, but only in 12 tissues (ps<0.05) (**Figure 6D**). Interestingly, in 13 tissues individuals with higher ISR activation exhibited lower mito-nDNA% (ps<0.05), demonstrating strong tissue specificity and a relatively equal number of tissues with positive and negative associations between ISR and mitochondrial gene expression. The strength and direction of the correlations between mitochondrial gene expression and either PGC-1α or the ISR were not significantly correlated (r=0.02-0.18, p=0.25-0.90), indicating that these pathways are not consistently co-regulated across human tissues.

Thus, if a nutritional, behavioral or other stressor activated PGC-1α or ISR systemically, and if these pathways influenced mitochondrial gene expression, we would expect that some tissues would exhibit an increase, some tissue would not change, while others would exhibit a decrease in mitochondrial gene expression and/or biogenesis in response to the same stimulus.

At the level of correlations, these organ-specific regulatory pathways would produce the lack of coherence and some apparent negative tradeoffs, as observed in our mouse and human cohorts.

### The discrepancy between mtDNA and nDNA-encoded transcripts is explained by tissue proliferation

The correlations between PGC-1α or ISR with mito-nDNA% were remarkably different than those with the marker of mitochondrial content, mtDNA% (**Supplemental Figure 8**). For example, in the heart (left ventricle), whereas PGC-1α expression was positively correlated with the fraction of the nuclear genome devoted to mitochondria as expected (mito-nDNA%, r=0.63, p<0.0001, **Figure 6A**), it was negatively correlated with the abundance of mtDNA-derived transcripts (mtDNA%, r=-0.36, p<0.0001, **Supplemental Figure 8A**). Brain tissues also exhibited this seemingly counterintuitive pattern where nDNA and mtDNA mitochondrial transcripts appear uncoupled or negatively correlated (**Figure 6A**). Negative correlations between mtDNA and nDNA transcripts have been reported previously in human brain tissues from the GTEx cohort ^29,30^. The same pattern of opposite associations across certain tissues was observed for the ISR pathway (see **Supplemental Figure 8** for the heart as an example), suggesting that a cell-level or tissue-level factor is responsible for these tissue differences in the coupling of mtDNA- and nDNA-derived transcripts.

Based on i) the dilution of mitochondria occurring during cell division in proliferating tissue (digestive tract, reproductive organs) ^31,32^, and ii) the striking stability of mitochondrial proteins in non-replicative tissues (e.g., brain, heart, muscle) ^33,34^, we reasoned that this discrepancy between mtDNA and nDNA-encoded transcripts and their associations with drivers of biogenesis could be attributable to tissue-specific differences in proliferative activity. Indeed, tissue proliferation (indexed by the average expression of 3 key proliferation genes: KI67, TOP2A, RRM2) ^35,36^ showed that the correlation of mtDNA% vs mito-nDNA% was positively associated with proliferation. In highly proliferative tissues (where newly made mitochondria are rapidly diluted by cell division; **Figure 7C**, bottom), tissues investing a large portion of their nuclear transcriptome into mitochondrial biogenesis also have high mtDNA-derived transcripts (positive correlation), likely as a means to replenish the ∼50% dilution that regularly takes place in symmetrically dividing cells. On the other hand, non-proliferative tissues with high mitochondrial mass tend to have proportionally lower nDNA-derived transcripts than the abundant mtDNA-derived transcripts contained in their numerous cytoplasmic mitochondria, consistent with the ability to maintain mitochondrial mass without sustained nuclear mitochondrial biogenesis in post-mitotic tissues (**Figure 7C**, top). Tissue replicative activity therefore accounts for the relative abundance of mtDNA- and nDNA-derived mitochondrial transcripts across human tissues.

**Figure 7.**
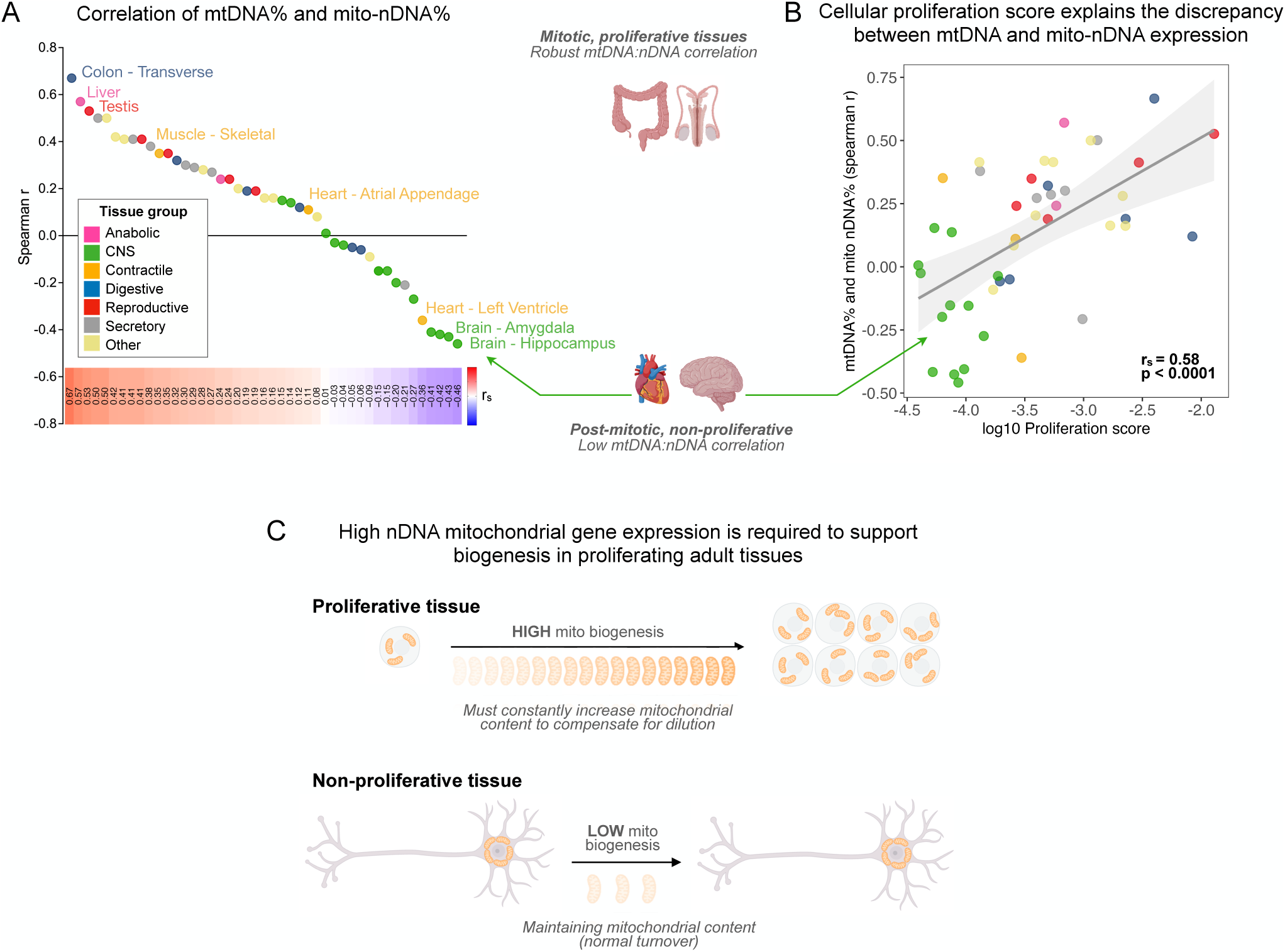
Discrepancy between mtDNA and mito-nDNA transcripts is explained by proliferation gene expression. (A) Ranked spearman r correlation coefficients of mtDNA% and mito-nDNA% transcripts in each tissue. (B) Biplot of proliferation score (average of KI67, TOP2A, RRM2) and spearman correlation coefficient of mtDNA% vs mito-nDNA% in each tissue. (C) Illustration of mitochondrial biogenesis in proliferative and non-proliferative tissues. In non-proliferative tissues that are also rich in mitochondria, such as heart and brain, transcripts for nDNA-encoded mitochondrial proteins are relatively low as biogenesis is not prioritized, nevertheless mtDNA-encoded transcripts remain high due to the abundance of mitochondria. However, in actively proliferating tissues, such as the colon and testes, mitochondrial biogenesis is upregulated resulting in high mito-nDNA and mtDNA transcripts in order to replenish the regular dilution of mitochondria in dividing cells.

### Sub-groups of individuals display distinct multi-tissue mitochondrial distribution patterns

Our analysis of the multi-tissue correlation structure in GTEx showed that individuals can have a high mitochondrial investment in some tissues, but low in other tissues. Based on the notion that different organs age at different rates in different individuals ^9–11^, this led us to examine if there are sub-groups or clusters of individuals who exhibit distinct multi-tissue mitochondrial distribution patterns. To address this question, we performed k-means clustering on mitochondrial gene expression data from 113 subjects with complete data across 4 tissues, interrogating the ratios in mito-nDNA% between each tissue pairs: Brain (cortex), Heart (atrial appendage), Muscle (skeletal), Adipose (subcutaneous) (**Figure 8A**).

**Figure 8.**
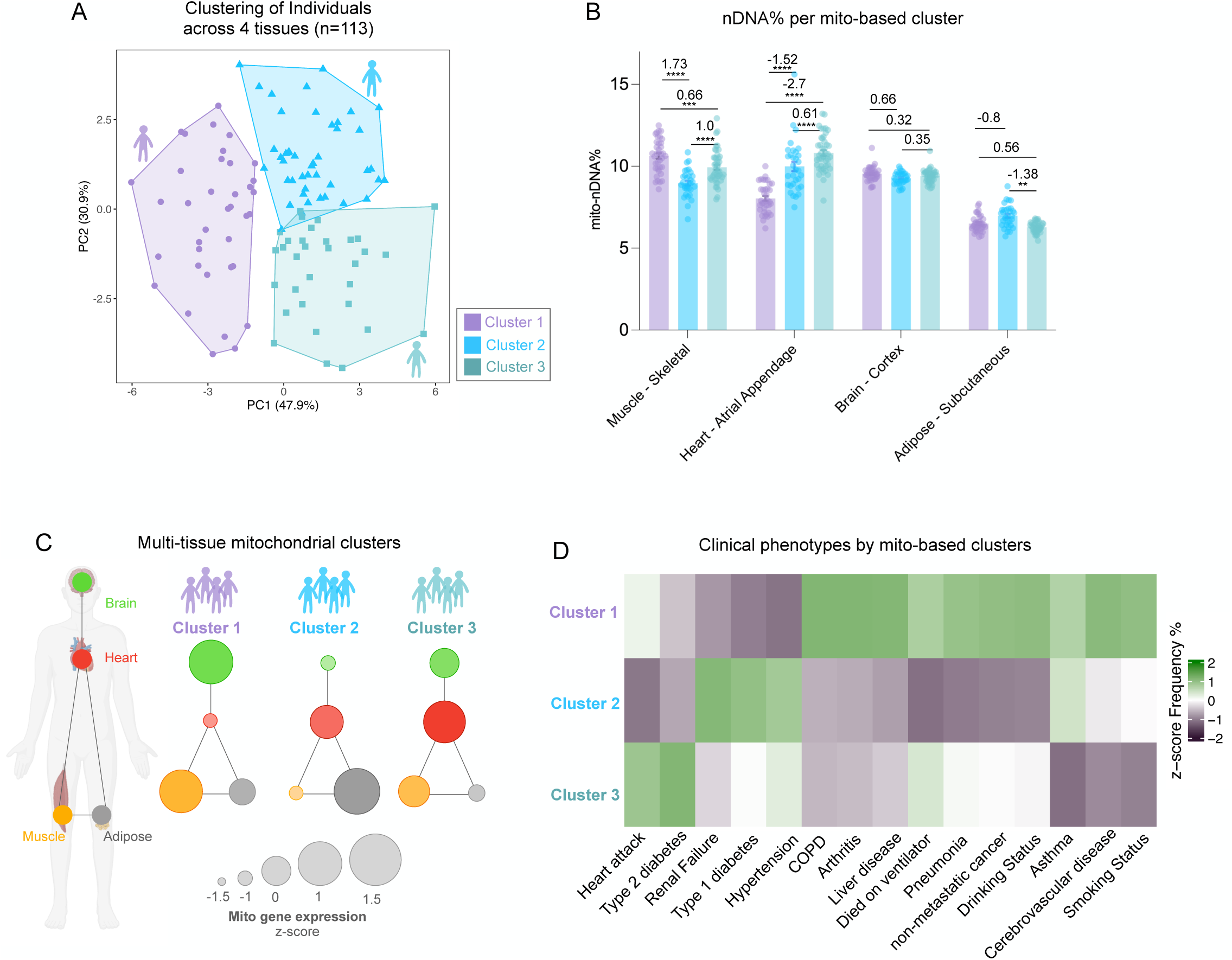
Sub-groups of individuals display different mitochondrial distribution patterns. (A) K-means clustering on mito-nDNA% ratios between 4 tissues (Muscle - skeletal, Heart - Atrial Appendage, Brain - Cortex, Adipose - Subcutaneous) from 113 subjects. Cluster 1 (n = 37), cluster 2 (n = 32), cluster 3 (n = 44). (B) Bar plot of mean mito-nDNA% in each cluster across 4 tissues. Cluster means of each tissue were tested for significant difference by two-way ANOVA. Effect sizes were computed by hedge’s g. (C) Network visualization of z-score transformed mito-nDNA% of each cluster across the 4 tissues analyzed. (D) Heatmap showing the z-score percentage of subjects in each cluster recorded as positive for each clinical variable.

Principal component analysis (PCA) of these data showed that the first two principal components accounted for 78.8% of variance, from which 3 clusters of GTEx participants emerged. A two-way ANOVA on the mito-nDNA% values for each tissue confirmed that the identified subgroups of individuals exhibited significant differences in mitochondrial transcript abundance across the 4 tissues analyzed (**Figure 8B-C**). The mitochondria-defined subgroups differed on clinical and phenotypic profiles including causes of death and known medical diagnoses at the time of death (**Figure 6D**). This association among multi-tissue mitochondrial profiles and clinical phenotypes provides preliminary evidence that distinct mitochondrial distribution strategies among the multi-organ network may be associated with different resilience or vulnerability profiles to disease.

## Discussion

Here we tested the hypothesis that the regulation of mitochondrial biology among dozens of organs and tissues is driven by an individual trait-level factor. We address this question by examining the multi-tissue correlation structure among direct enzymatic measurements of mitochondrial OxPhos respiratory chain enzyme activities in two cohorts of mice, and from the transcriptome and mtDNAcn of 45 human tissues. Contrary to the default hypothesis that mitochondrial content is a trait-like attribute of each person exhibiting high coherence across the organism, our results show a striking spectrum of mitochondrial distribution patterns. In particular, we show that individuals who have high markers of mitochondrial content in some tissues can have low markers of mitochondria in other tissues. We find that these differences may be partially explained by the energy sensing pathways PGC-1α and ISR, and by proliferative activity of different tissues, expanding our understanding of the factors that may regulate the highly heterogenous and tissue specific regulation of mitochondrial biology in humans and other animals. These results emphasize how genetic and systemic factors are insufficient to explain the mitochondrial expression patterns across the brain and body, suggesting instead that different individuals display relatively idiosyncratic patterns of mitochondrial regulation.

The results of our investigation of the inter-tissue correlation structure of mitochondria in mouse and human tissue is consistent with there being partial coherence between the organs/tissues of an individual. This indicates that there is a certain degree of inter-tissue coregulation of mitochondria, particularly between tissues of the same type. We observed distinct distributions in coherence between brain-brain, body-body, and brain-body tissues.

Different brain regions showed the highest coherence. However, there was minimal evidence of coherence between brain and body tissues. The specific tissues with the highest average coherence were esophagus - gastroesophageal junction and esophagus - muscularis. Adrenal gland, liver and kidney displayed the lowest average inter-tissue coherence. Overall, given that the highest correlations of mito-nDNA% did not exceed r=0.6, and many tissues show no evidence of coherence, this indicates that every person may have an idiosyncratic mitochondrial distribution pattern, contributing to the uniqueness of each individual.

Our analysis of the relationship between mitochondrial gene expression and energetic stress sensing metabolic pathways (PGC-1α and ISR) revealed a strongly tissue-dependent association. The results for PGC-1α were particularly striking. While consistent with previous evidence in skeletal muscle and heart, it appears that PGC-1α does not drive mitochondrial biogenesis in all tissues at the transcript level. In fact, an opposite association was observed in several tissues, calling for a more refined understanding of the forces that regulate mitochondrial biology and biogenesis in particular, in different organ systems. The novel association of mtDNA:nDNA ratio with proliferative activity across tissues provides an example of such a force that governs mitochondrial biogenesis in a tissue-specific manner.

The significant differences in mitochondrial gene expression patterns between subgroups of individuals suggest that some individuals “invest” more of their energetic resources in some tissues compared to others. This aligns with recent studies identifying distinct ageotypes (i.e., aging phenotypes or strategies) among individuals ^9,11,37^. In our data, these groups also displayed different patterns of clinical phenotypes, opening the possibility that distinct mitochondrial distribution patterns among the multi-organ network may be associated with differences in resilience or vulnerability to specific diseases. This notion aligns with an emerging systemic understanding of mitochondrial biology with regulatory signals and even whole mitochondria transferred between organ systems and cell types ^2,38^. It remains an open question whether these clusters are permanent traits of an individual (determined, for example, by genetics or developmental conditions), or are adjusted temporally within an individual based on current conditions.

### Study limitations

An important limitation of this study is the use of post-mortem tissue. In GTEx, post-mortem interval time is associated with tissue-specific changes in transcripts abundance ^39^. This could have affected the transcriptome-based mitochondrial profiling and inter-tissue correlation analyses using GTEx data, but not the mtDNAcn-based analyses in human tissues nor the enzymatic and mtDNA-based measures in mouse tissue. Moreover, our sensitivity analyses showed that our transcriptome-based inter-tissue correlation patterns were stable across various RNA integrity number (RIN) cut-offs. Another limitation is that not all tissues were available for all individuals, which limited the number of tissue-tissue pairs available for some pairs of organs and tissues. Future studies with complete data in greater numbers of tissues and individuals, and with direct measures of mitochondrial functions (respiration, OxPhos, ROS production), would be useful in extending and validating these results.

## Methods

### Mouse tissue homogenization, enzymatic activity assays and qPCR

Enzymatic activity assays and qPCR were performed on mouse tissue as described previously ^40^. The first mouse cohort included n=16 c57bl/6J female and male mice, from which 5 tissues were sampled, as described in ^41^. The second mouse cohort (n=27 mice, 22 tissues) is described in ^40^. All cohort 2 mice were 52-week-old male c57bl/6J animals. mtDNA density was quantified as described in ^40^. Raw data was used to compute inter-tissue correlations for each mitochondrial feature including Complex I activity, Complex II activity, Complex IV activity, Citrate Synthase activity and mtDNA density. A total of 50 (10 per mitochondrial feature) comparisons were computed for Cohort 1, and 1,155 (231 per mitochondrial feature) for Cohort 2.

### GTEx dataset

The GTEx v8 RNAseq dataset consists of 17,382 samples across 54 tissues from 948 donors. The GTEx cohort is 67.1% male and 32.9% female, among which 84.6% are White, 12.9% are African American, 1.3% are Asian and 1.1% other. The age range of donors is 20-70 years (mean ± SD = 52.8 ± 12.9). Further information on the GTEx v8 dataset can be found on the GTEx portal at https://gtexportal.org/home/tissueSummaryPage.

### Transcriptomics

RNA sequencing data was downloaded as TPM from the GTEx portal. Data is available for download at https://gtexportal.org/home/downloads/adult-gtex/bulk_tissue_expression.

Subject clinical phenotype data is part of the protected access data and was obtained through the dbGAP accession #phs000424.v8.p2 under project #27813 (Defining conserved age-related gene expression trajectories). All samples with an RNA integrity number (RIN) of less than 5.5 were filtered out of the data. For inter-tissue correlation analysis, a minimum sample size of 10 shared subjects was set for each tissue pair and all tissue pairs with less than 10 subjects were not included in the analysis. Tissues that did not meet this shared sample size with more than 50% of other tissues were not included in the analysis. Cell lines were also removed from the dataset. After applying these filters to the data, 45 tissues and 16,205 samples remained and were included in the analysis. The mitochondrial genes included in analyses were the Mitocarta 3.0 genes (of which we identified 1133 in the dataset) and all other mtDNA genes. TPM values were normalized separately for mtDNA and nDNA genes. mtDNA genes were expressed as a percentage of all transcripts in a sample. Nuclear genes were normalized by removing all mtDNA transcripts from the sample, and then expressed as a percentage of all remaining transcripts (proportion of nuclear transcriptome). We performed a sensitivity analysis by repeating our mito-nDNA% inter-tissue correlation analysis with a sample RIN cut-off of RIN>6 and RIN>7. This had a negligible effect on the median inter-tissue correlation and correlation structure, but it greatly reduced the number of tissue pairs that could be included in the analysis. Therefore, we chose a less stringent RIN cut-off of 5.5 to maximize inter-tissue sample sizes and the number of tissue pairs included in our analysis.

### Proteomics

Proteomics data were obtained from the supplemental information in Jiang et al 2020 ^19^. Supplemental table S2C “protein normalized abundance” was downloaded and used for analysis. The abundances of mitochondrial genes, of which we identified 1002, were summed and expressed as a percentage of total protein abundance to establish a proximate marker of mitochondrial abundance in each sample. Replicate measurements of samples from multiple proteomics runs were averaged for the pairwise inter-tissue correlation analysis. Only tissue pairs with a minimum of 8 shared subjects were included in the analysis. Tissues that did not meet this minimum sample size with at least 4 other tissues were excluded. This resulted in 7 tissues and 14 subjects included in the analysis.

### mtDNAcn

mtDNAcn data on the GTEx cohort were downloaded from the supporting information in Rath et al 2024 ^15^. The data was filtered for the same 45 tissues used in transcriptomics data analysis. Only tissue pairs with a minimum of 10 shared subjects were included in analysis. Individual mtCN values for each GTEx tissue sample were used for inter-tissue correlation analysis.

### Tissue proliferation index

Our criteria for selecting genes to include in the proliferation index were as follows; (i) the genes are known in the literature to be markers of proliferation ^35,36^, (ii) the gene must display high expression in tissues that are known to be proliferative (e.g. blood, digestive tract) ^31^, (iii) the genes must show internal consistency (i.e. positively correlated within tissues), (iv) the genes must display high expression during periods of rapid cell division in a human primary tissue culture system ^42^. The genes selected for the tissue proliferation index were KI67, TOP2A, RRM2. The index was established by averaging the expression of these 3 genes in each tissue sample.

### Multi-tissue Network Graph

The multi-tissue network graph in figure 5 was generated with the R package iGraph. An adjacency matrix was generated from the inter-tissue correlation matrix shown in Figure 4B, setting the edge threshold to r=0.2. The layout of nodes was determined by Fruchterman-Reingold layout force-directed algorithm.

### Statistics

Statistical tests were performed with R version 4.4.0 (2024-04-24) and GraphPad Prism version 10. The correlations between pairwise tissue comparisons were assessed using spearman rank correlation. Clustering was performed on inter-tissue ratios using k-means method and was visualized using principal component analysis. Two-way ANOVA was used to test mitochondrial gene expression of identified clusters for significant differences. Effect sizes were estimated using hedge’s g. The significance level for P-values was set at P<0.05.

## Code and data availability

The GTEx v8 RNAseq dataset can be downloaded from the GTEx portal at https://gtexportal.org/home/downloads/adult-gtex/bulk_tissue_expression. All R code used in the analyses can be found at github.com/mitopsychobio.

## Acknowledgements

Portions of figures were created with Biorender.com.

## Author contributions

M.P. and A.A.C conceived the project. A.J. and J.D. collected data on mouse cohort 1.

A.M.R collected data on mouse cohort 2. J.D. and A.S.M. performed the analyses. J.D generated the figures. M.P., J.D., A.A.C, A.S.M, and D.S. interpreted the data. J.D. and M.P. drafted the manuscript. All authors reviewed and edited the final manuscript.

## Supplemental Material

**Supplemental File 1.** Excel file containing the two mouse mitochondrial enzymatic activity and mtDNA density datasets used in analyses. Sheet 1 contains mouse cohort 1 dataset. Sheet 2 contains mouse cohort 2 dataset.

**Supplemental File 2.** Excel file containing the average, standard deviation, minimum and maximum values of mito-nDNA% (sheet 1) and mtDNA% (sheet 2) in each of the 45 tissues analyzed using the GTEx RNAseq v8 dataset.

**Supplemental Figure 1.** (A) Correlation matrix of enzymatic activity measures (CI, CII, CIV and CS) and mtDNA density across 5 tissues (hippocampus, brown fat, liver, muscle and bone) from 16 male mice in Cohort 1. (B) Frequency distribution of spearman r correlation coefficients in which only the inter-tissue correlations of the same measure are included (n=50 pairwise tissue comparisons). (C-E) Bivariate plots of mitochondrial enzyme activity measures between tissues.

**Supplemental Figure 2.** Correlation of average mtDNA% with average mito-nDNA% across 45 tissues.

**Supplemental figure 3.** Heatmap showing sample size of every pairwise tissue comparison in the same order as Figure 4B.

**Supplemental figure 4. Multi-tissue mitochondrial correlation patterns of mitochondrial pathways.** Correlation matrix and frequency distribution of correlation coefficients of mtDNA genes, OxPhos genes, fission and fusion genes, ROS and glutathione metabolism.

**Supplemental Figure 5. Multi-tissue network architecture of mitochondrial gene expression based on mtDNA-encoded genes**. Network representation of multi-tissue correlation of mtDNA%. Each node represents a tissue, the size of the node is proportional its degree, edge thickness is proportional to the strength of the correlation.

**Supplemental Figure 6. mtDNAcn-based inter-tissue correlation structure.** (A) Heatmap of correlation matrix showing the pairwise spearman r correlations of mtDNAcn between 45 tissues. (B) Frequency distribution of spearman r correlation coefficients between brain-brain (red), body-body (green) and brain-body (yellow) tissues. (C) Network architecture of mtDNAcn-based inter-tissue correlations. (D-F) Bivariate plots of mtDNAcn between tissues.

**Supplemental Figure 7. Proteomics-based mitochondrial correlation patterns across 7 tissues from 14 GTEx subjects** ^19^. (A) Heatmap of correlation matrix showing pairwise tissue comparisons of mitochondrial protein abundance. (B) Frequency distribution of spearman correlation coefficients of inter-tissue correlations of mitochondrial protein abundance. (C-E) Bivariate plots of mitochondrial protein abundance between tissues.

**Supplemental Figure 8. Mitochondrial gene expression is driven in part by canonical energy and stress sensing metabolic pathways.** (A) Ranked spearman r correlation coefficients of PGC1a% vs mtDNA% transcripts in each tissue. (B) Biplot showing the correlation of mtDNA% with PGC-1α expression in the colon (transverse). (C) Biplot showing the correlation of mtDNA% with PGC-1α expression in the anterior cingulate cortex. (D) Ranked spearman r correlation coefficients of ISR% vs mtDNA% transcripts in each tissue. (E) Biplot showing the correlation of mtDNA% with ISR expression in the heart (left ventricle). (F) Biplot showing the correlation of mtDNA% with ISR expression in the colon (transverse).

**Supplemental Figure 9. Sub-groups of individuals display different mitochondrial distribution patterns.** (A) K-means clustering on mtDNA% ratios between 4 tissues (Muscle - skeletal, Heart - Atrial Appendage, Brain - Cortex, Adipose - Subcutaneous) from 113 subjects. Cluster 1 (n = 46), cluster 2 (n = 29), cluster 3 (n = 38). (B) Bar plot of mean mtDNA% in each cluster across 4 tissues. Cluster means of each the tissue were tested for significant difference by two-way ANOVA. Effect sizes were computed by hedge’s g. (C) Network visualization of z-score transformed mito-nDNA% of each cluster across the 4 tissues analyzed. (D) Heatmap showing the z-score percentage of subjects in each cluster who are recorded as positive for each clinical variable.

