## Supplemental Figures for "Brain-body mitochondrial distribution patterns lack coherence and point to tissue-specific and individualized regulatory mechanisms"

### Supplemental Figure 1

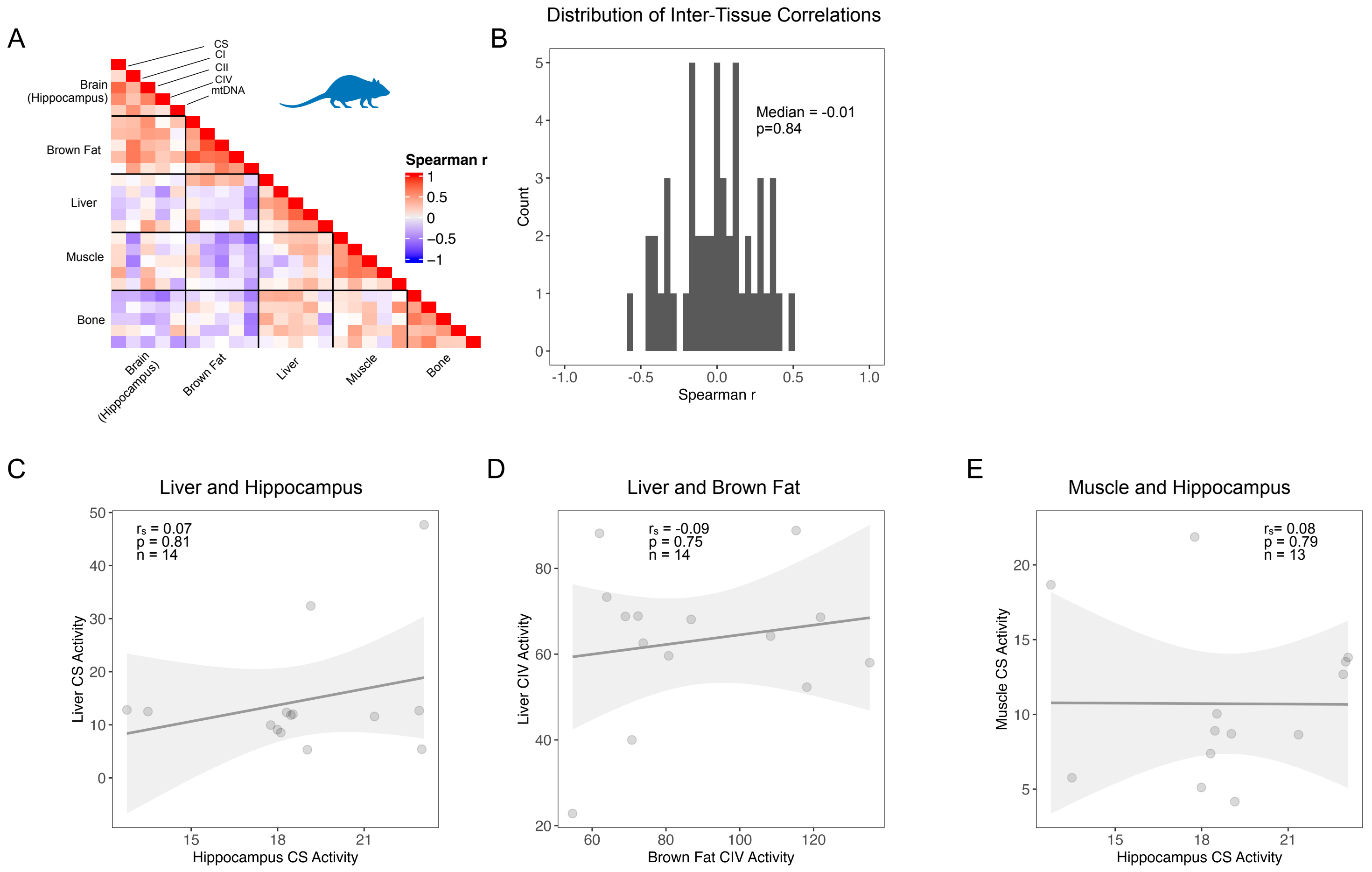

### Supplemental Figure 2

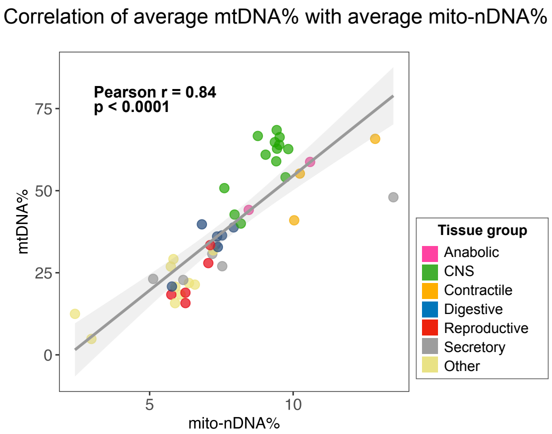

### Supplemental Figure 3

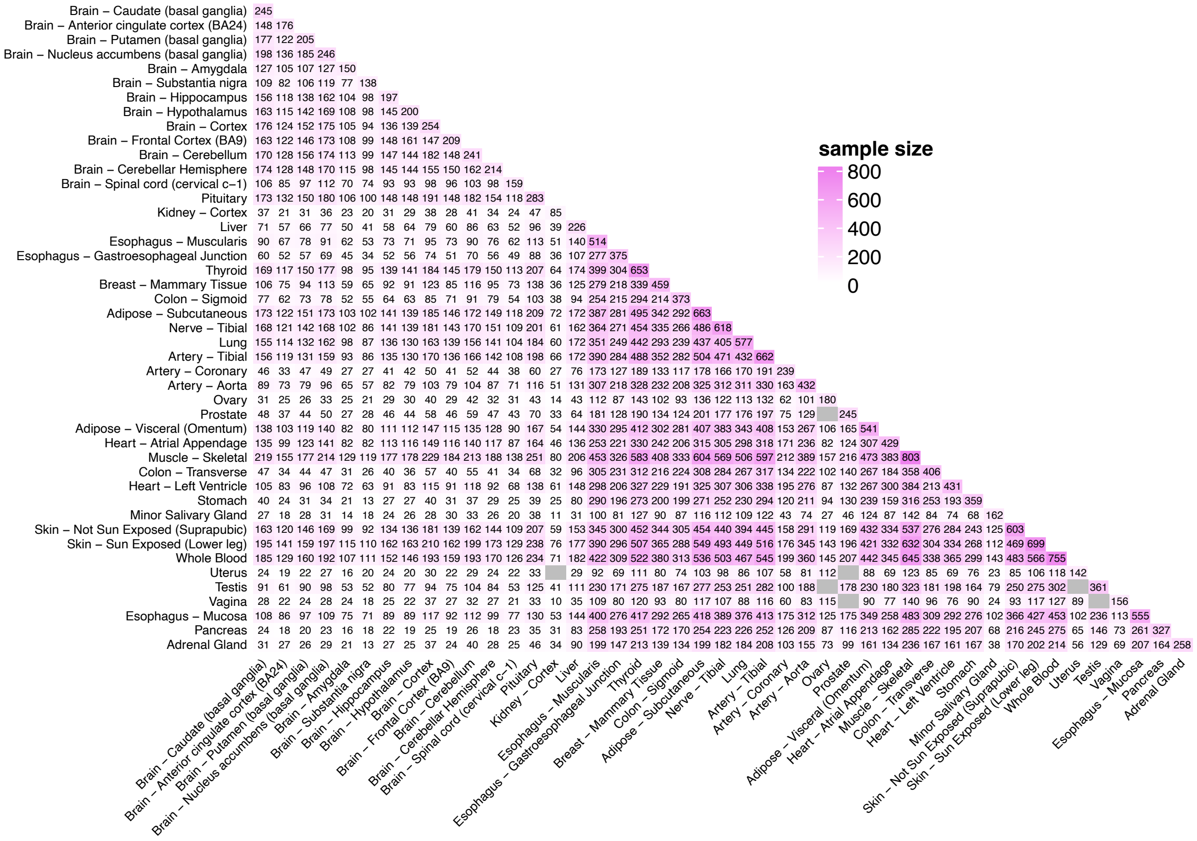

#### Supplemental Figure 4

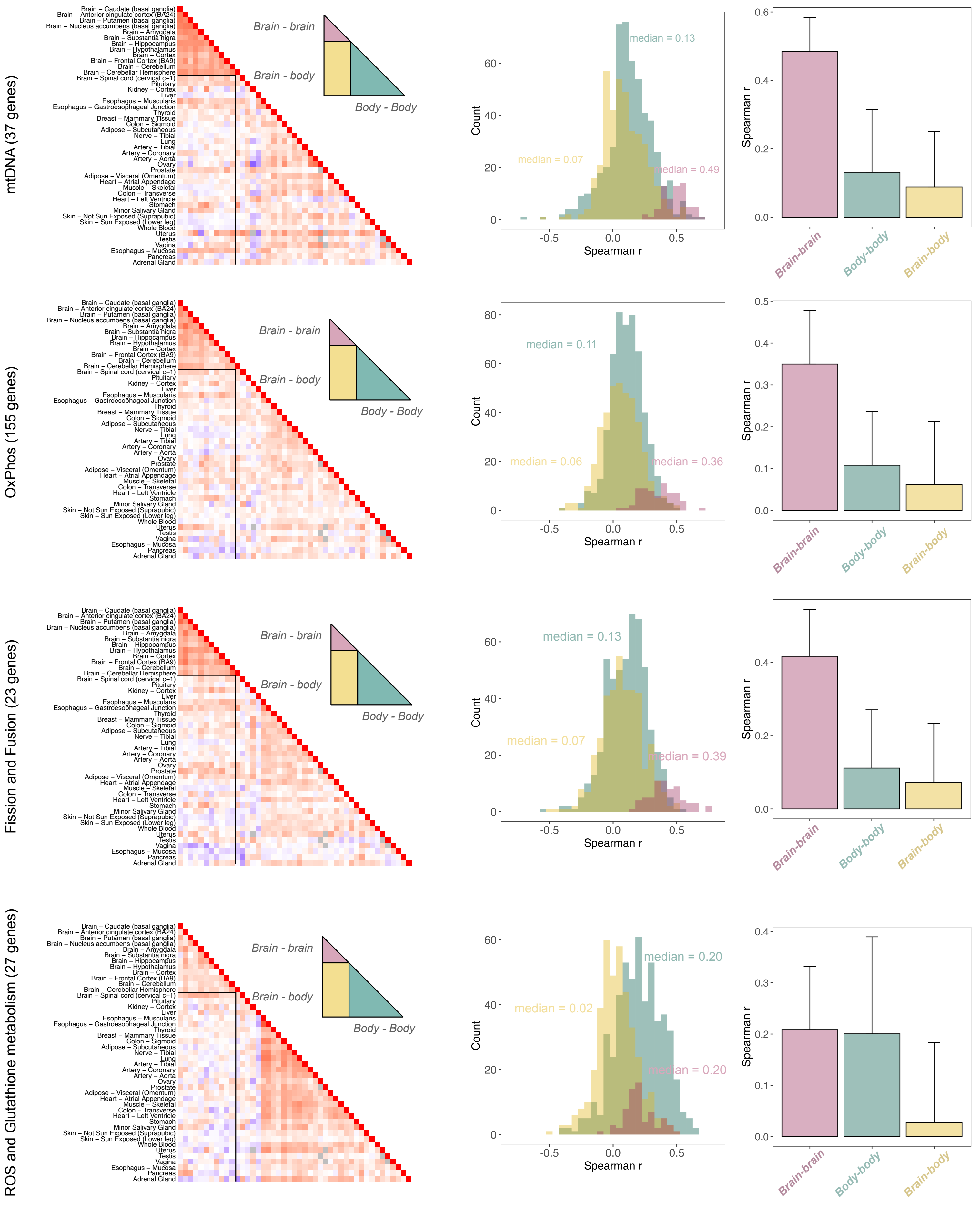

### Supplemental Figure 5

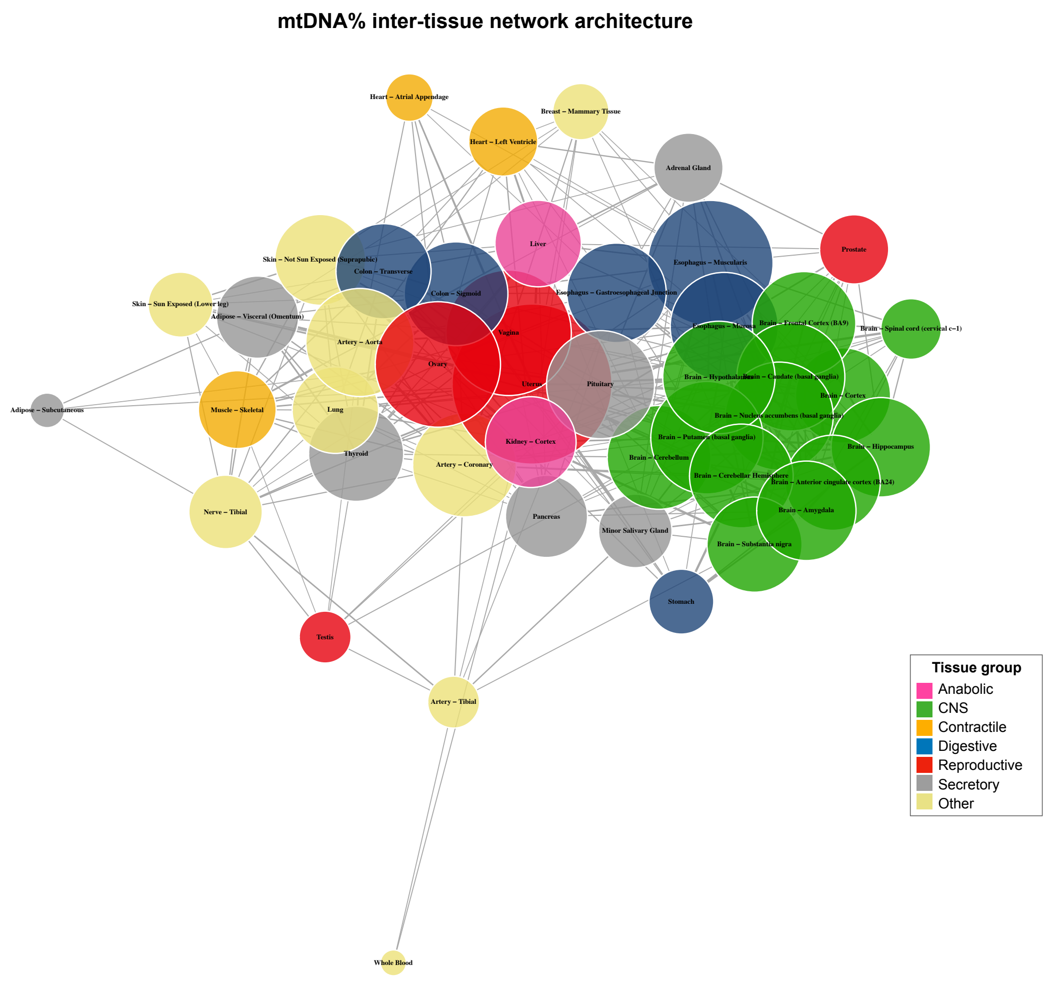

### Supplemental Figure 6

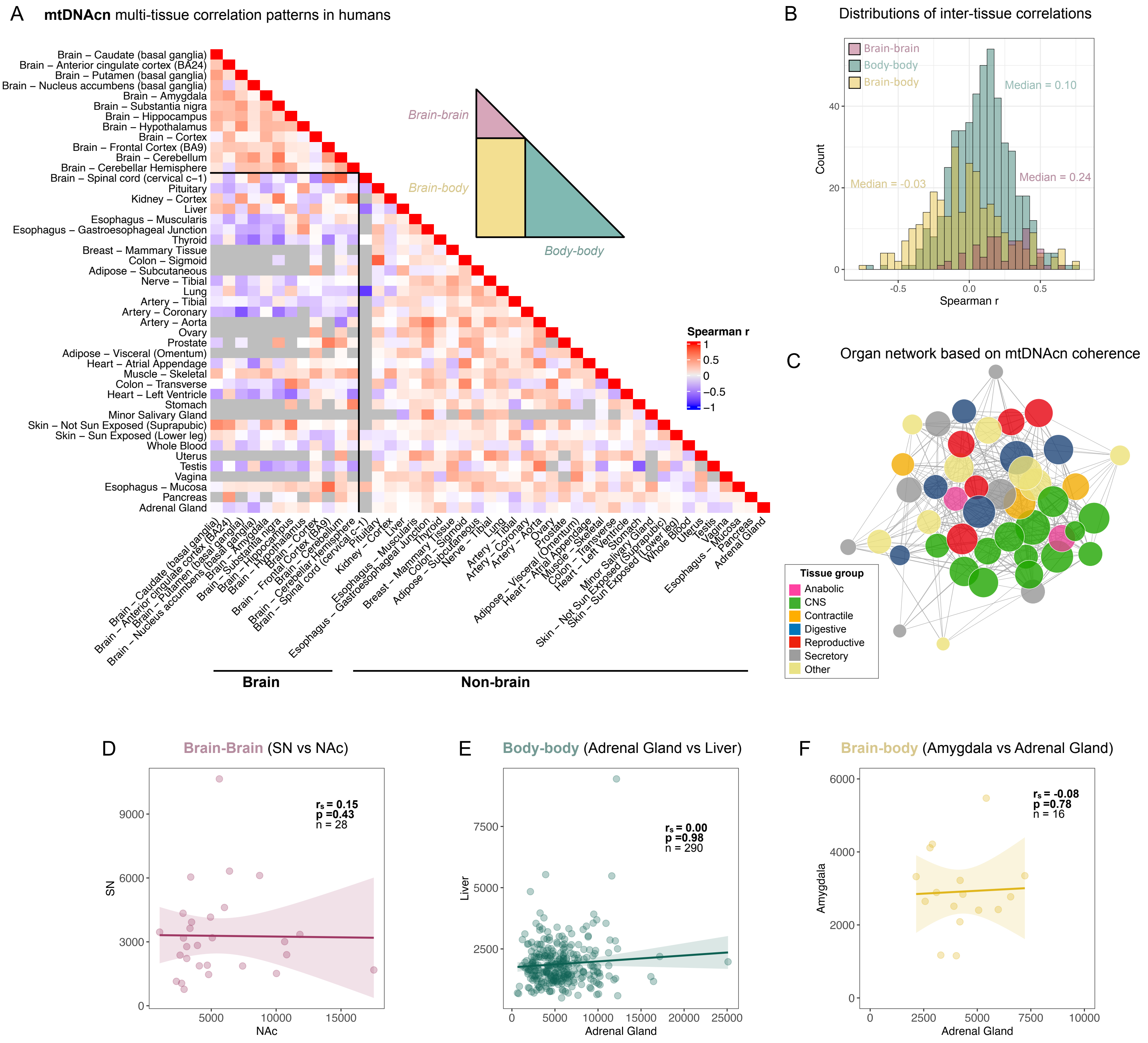

### Supplemental Figure 7

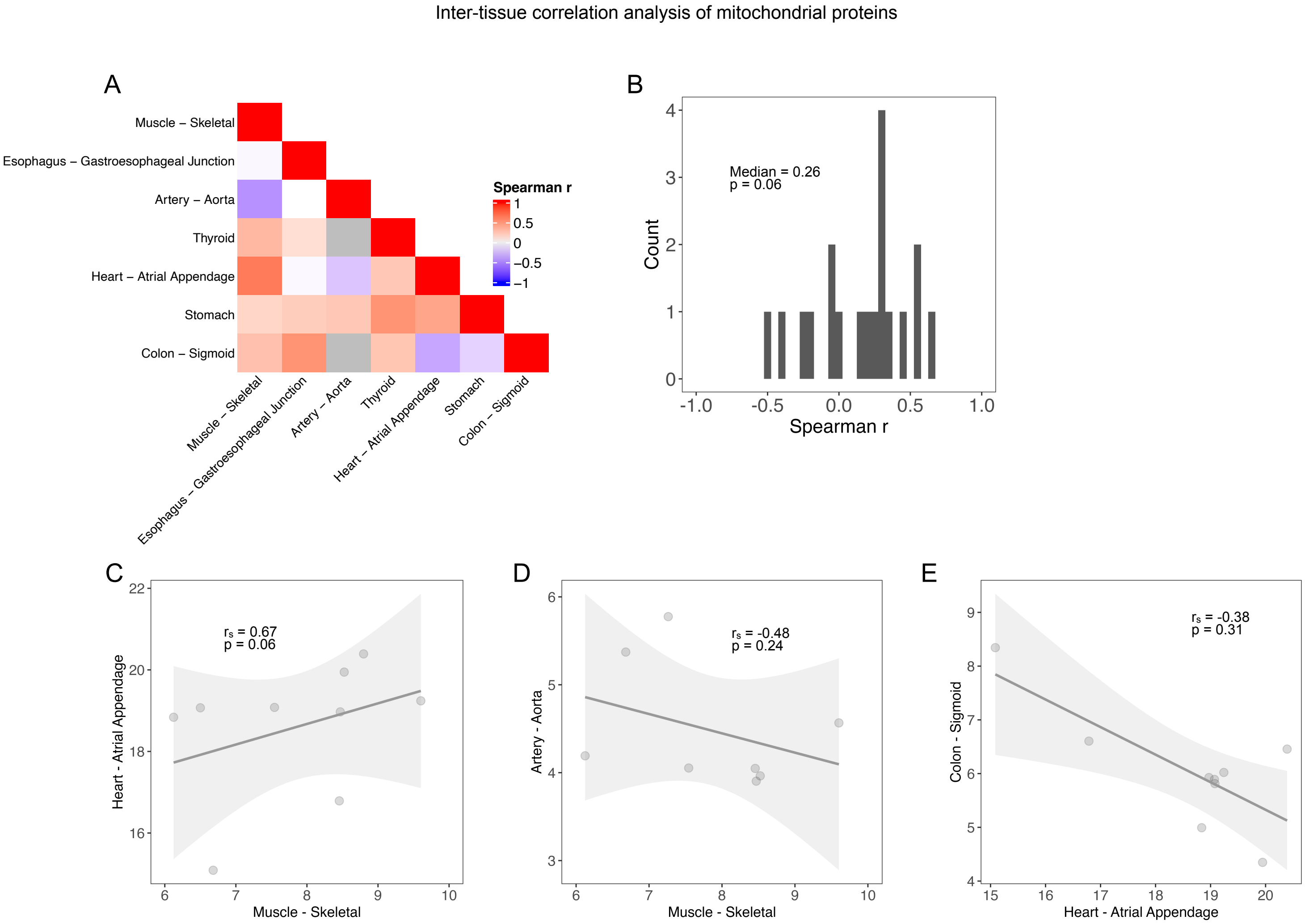

### Supplemental Figure 8

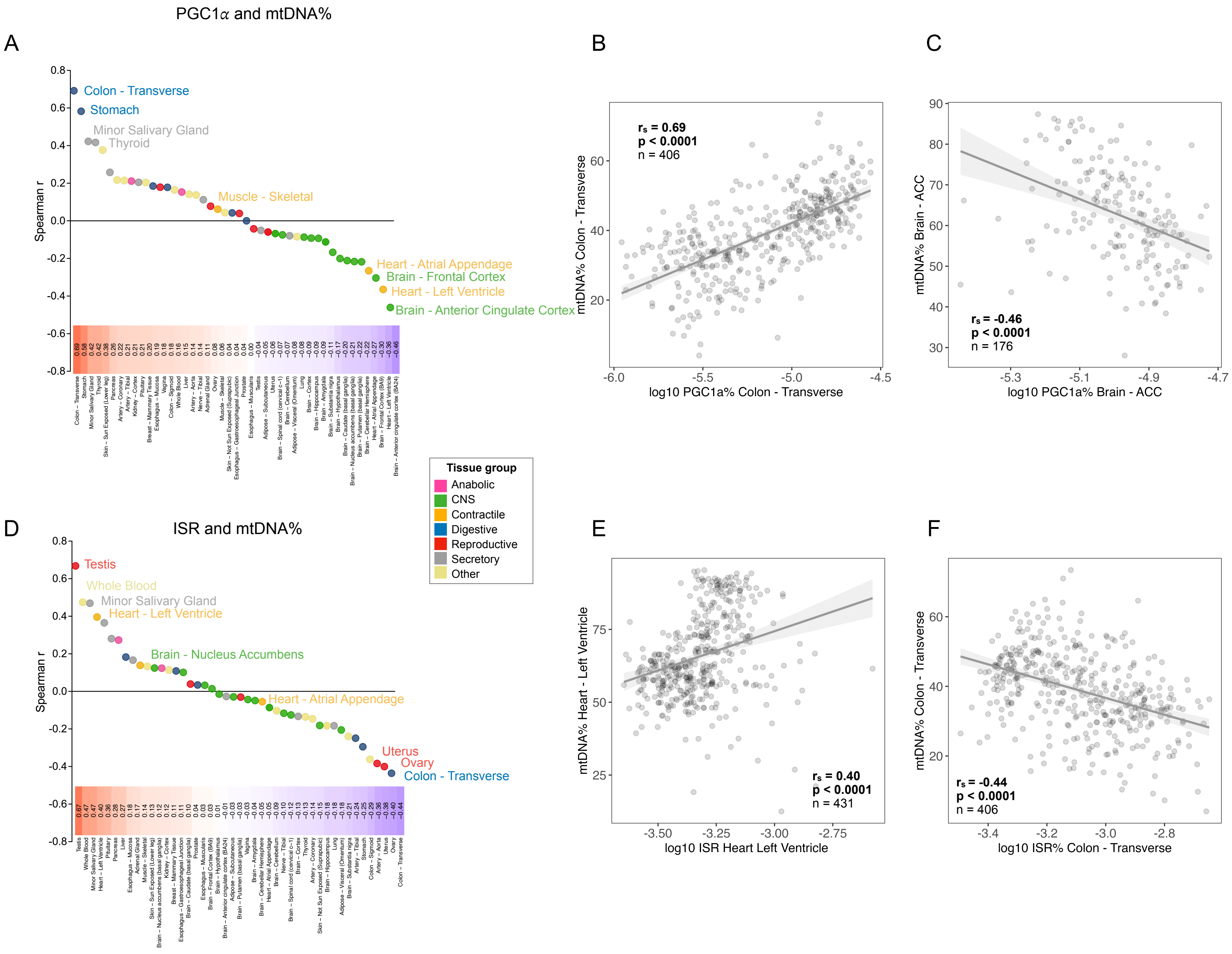

### Supplemental Figure 9

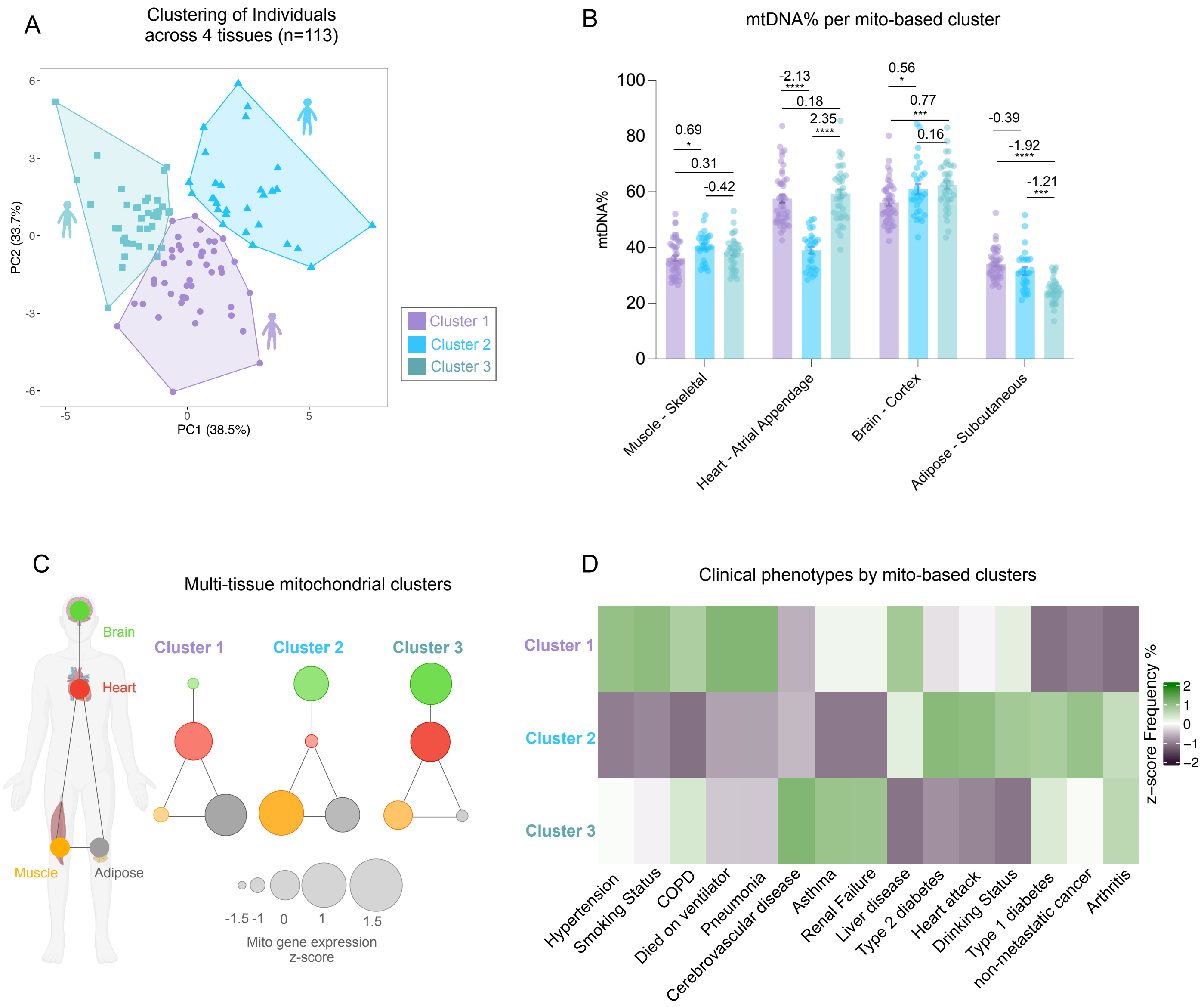
